# Beyond planar: fish schools adopt ladder formations in 3D

**DOI:** 10.1101/2024.10.03.616549

**Authors:** Hungtang Ko, Abigail Girma, Yangfan Zhang, Yu Pan, George V Lauder, Radhika Nagpal

## Abstract

The coordinated movement of fish schools has long captivated researchers studying animal collective behavior. Classical literature from Weihs and Lighthill suggests that fish schools should favor planar diamond formations to increase hydrodynamic efficiency, inspiring a large body of work ranging from fluid simulations to hydrofoil experiments. However, whether fish schools actually adopt and maintain this idealized formation remains debated and unresolved. When fish schools are free to self-organize in three dimensions, what formations do they prefer? By tracking polarized schools of giant danios (*Devario aequipinnatus*) swimming continuously for ten hours, we demonstrate that fish rarely stay in a horizontal plane, and even more rarely, in the classical diamond formation. Of all fish pairs within four body-lengths from each other, only 25.2% are in the same plane. Of these, 54.6% are inline, 30.0% are staggered, and 15.4% are side-by-side. The diamond formation was observed in less than 0.1% of all frames. Notably, a “ladder formation” emerged as the most probable formation for schooling giant danios, appearing in 79% of all fish pairs and fish schools elongate at higher swimming speeds. These findings highlight the dynamic and three-dimensional nature of fish schools and suggest that hydrodynamic benefits may be obtained without requiring fixed positions. This research provides a foundation for future studies that examine the hydrodynamics and control of underwater collectives in 3D formations.

## 1 Introduction

Schooling by fish species is a common and widely studied canonical collective behavior. It has been estimated that 50% of fish species form schools as juveniles and 25% form schools throughout their life [1]. Recent experimental studies have demonstrated that schooling behavior allows fish to save up to 53% of metabolic energy compared to swimming by themselves [2]. One potential cause of this energetic saving is the complex hydrodynamic interaction between fish and their surrounding fluid environment, an intriguing problem that dates back to the 1970s, when applied mathematicians Weihs [3] and Lighthill [4] theorized that a planar diamond formation is both energy-saving and self-stabilizing. Using the potential flow theory in fluid mechanics, they argued that this formation would allow a trailing fish to avoid the jet produced by its leaders and swim in a reduced flow region. Furthermore, assuming synchronized kinematics, the side-by-side fish pairs in this formation would have an increased hydrodynamic efficiency.

While the diamond formation has inspired a diversity of work using simulations, hydrofoils, and underwater robots [5], whether fish schools adopt this configuration remains highly debated [6]. Recently, other planar formations have been identified in fish experiments [7, 8, 9, 10, 11, 12, 13, 14] (Table 1). For example, [10] showed that fish prefer a side-by-side ‘phalanx’ formation at a swimming speed of around 4 body-lengths per second. [12] demonstrated that rainbowfish (*Melanotaenia*) favor inline formations whose spacing increases with the swimming speed.

**Table 1.** A summary of selected experimental studies that directly characterize fish school formations both in two and three dimensions, and three key parameters relevant to the experimental arrangement used in this study.

|  | Presence of flow | Recording length | Water height |
| --- | --- | --- | --- |
| Butail & Paley 2012 | X | 3 s | 8 BL |
| Herbert-read et al 2011 | X | 5 min | 1 BL |
| Kent et al 2019 | X | 5 min | 1.2-1.8 BL |
| Mekdara et al 2018 2021 | X | 5 min | 12 BL |
| Katz et al 2011 | X | ~1 hr | 1 BL |
| Escobedo et al 2020 | X | ~1 hr | 1.9 BL |
| Ashraf et al 2016, 2017 | ✓ | 10 s | 0.5 BL |
| Lombana & Porfiri 2022 | ✓ | 5 min | 2.8 BL |
| **Current work** | ✓ | **10 hr** | **1.9 BL** |

Studying fish schooling behavior in laboratory settings presents significant challenges, particularly when attempting to identify the three-dimensional positions of multiple individuals swimming freely in deep water and against currents. Consequently, most experimental studies of fish school formations have been limited, often conducted either in still water, using short video recordings, or in shallow water to restrict vertical movement (Table 1). Without flow, it is difficult to distinguish fish positions resulting from social cohesive interactions or food-seeking behavior from those that lead to energetic savings – a consideration that becomes increasingly important as swimming speed increases [2]. To study fish school dynamics under active directional locomotion and identify positions that may offer beneficial hydrodynamic interactions, it is essential to provide a three-dimensional experimental condition where fish groups are challenged to swim actively in a common mean direction. In the absence of continuous flow, fish groups in still water tend to form non-polarized formations [15, 2], often swimming intermittently with varied headings. Such conditions are unsuitable for characterizing fish school formations that provide hydrodynamic benefits or for testing classical diamond formation proposed by Weihs and Lighthill [4, 3].

Previous studies on fish school behavior at higher swimming speeds often limit their recordings to less than five minutes [11, 14]. It is unclear whether this provides sufficient time for schools to reach a steady formation in a laboratory setting and if school kinematics might change over an extended duration. Do fish schools maintain consistent formations over hours when swimming against the flow? In this study, we address this question by recording active directional schooling behavior in small schools of giant danio (*Devario aequipinnatus*) for ten hours, generating long-duration 3-D datasets that enable comprehensive analysis of fish school formation dynamics. To ensure that fish can swim continuously for such a long period, we used a moderate speed at which fish schools exhibit the minimum cost of transport [2], at approximately 2.4 body-lengths per second.

Most previous experiments also confined fish schools to shallow water to facilitate automated tracking and individual identification (Table 1). This artificial constraint forces fish schools into planar formations near surface boundaries that can significantly substantially alter flow and influence group behavior. In natural environments such as rivers and ocean, however, fish schools exhibit threedimensional structures when provided sufficient water depth (Fig. 1). While a pair of fish may appear to be inline or staggered in a ventral view, they are often separated vertically. Such three-dimensional arrangements challenge the planar assumption fundamental for the diamond formation conjecture [3]. However, due to the difficulties associated with automatic tracking, long-term quantitative characterization of fish school formations in three dimensions remains limited.

**Figure 1.**
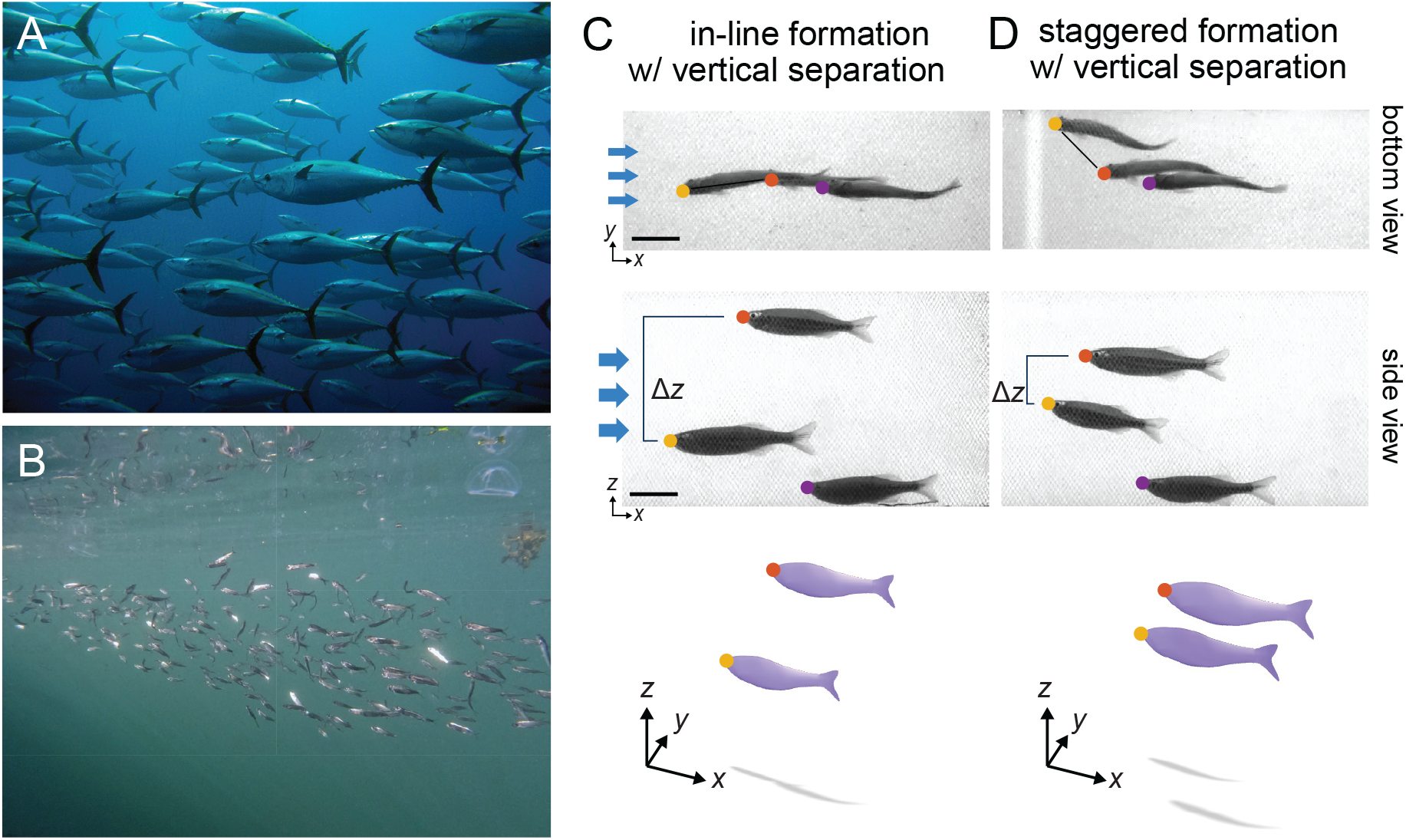
Fish schools adopt 3D formations both in nature and in the laboratory. Schools of (A) bluefin tuna and (B) pacific herring in 3D formations. (C-D) A small school of three giant danios with a pair (red and yellow dots) in an apparent (C) inline and (D) staggered formation. Both had a significant vertical separation.

Therefore, the goal of this study was to characterize the frequency of different three-dimensional fish formations within small experimental schools in the laboratory. We hypothesized that the 3D nature of fish schools could result in preferred formations that are non-planar, potentially challenging the assumptions of many common kinematic and hydrodynamic models used to support energy-saving resulting from specific formations. To test this, we conducted long-duration 3D recordings of small schools of giant danios (*Devario aequipinnatus*) swimming at 1.6 to 5.6 BL/s in a circulating flow channel. We developed a custom computer vision pipeline utilizing deep neural networks to extract the 3D coordinates of all fish in the schools. This combined approach allowed us to directly test the diamond formation conjecture and explore the preferred formations of giant danio in three dimensions.

## 2 Results

We placed schools of six giant danio in a chamber 8.8 times the height of a fish (1.9 BL)(Fig. 2A-B). Bounded by a mesh net and located within the middle of a recirculating flow tank, the chamber minimizes the effects of fluid boundary layers, as verified by particle imagery velocimetry (PIV) (Fig. S1). A flow of 2.4 body-lengths per second (BL/s) polarized the schools, reducing pitch *α* and yaw *ϕ* angles to near zero (Fig. 2C). The fish schools were recorded from both side and bottom views. For each view, we trained two neural network models using different computational architectures in SLEAP.ai [16] to locate the coordinates of each fish’s nose (Fig. 2D). A custom MATLAB program then integrated data from both models, removed outliers, triangulated 3D coordinates, and tracked individuals across frames over time (see methods and supplemental materials). Fig. 2E illustrates sample 3D trajectories of six fish over a two-minute period, highlighting the detailed behavioral data collected.

**Figure 2.**
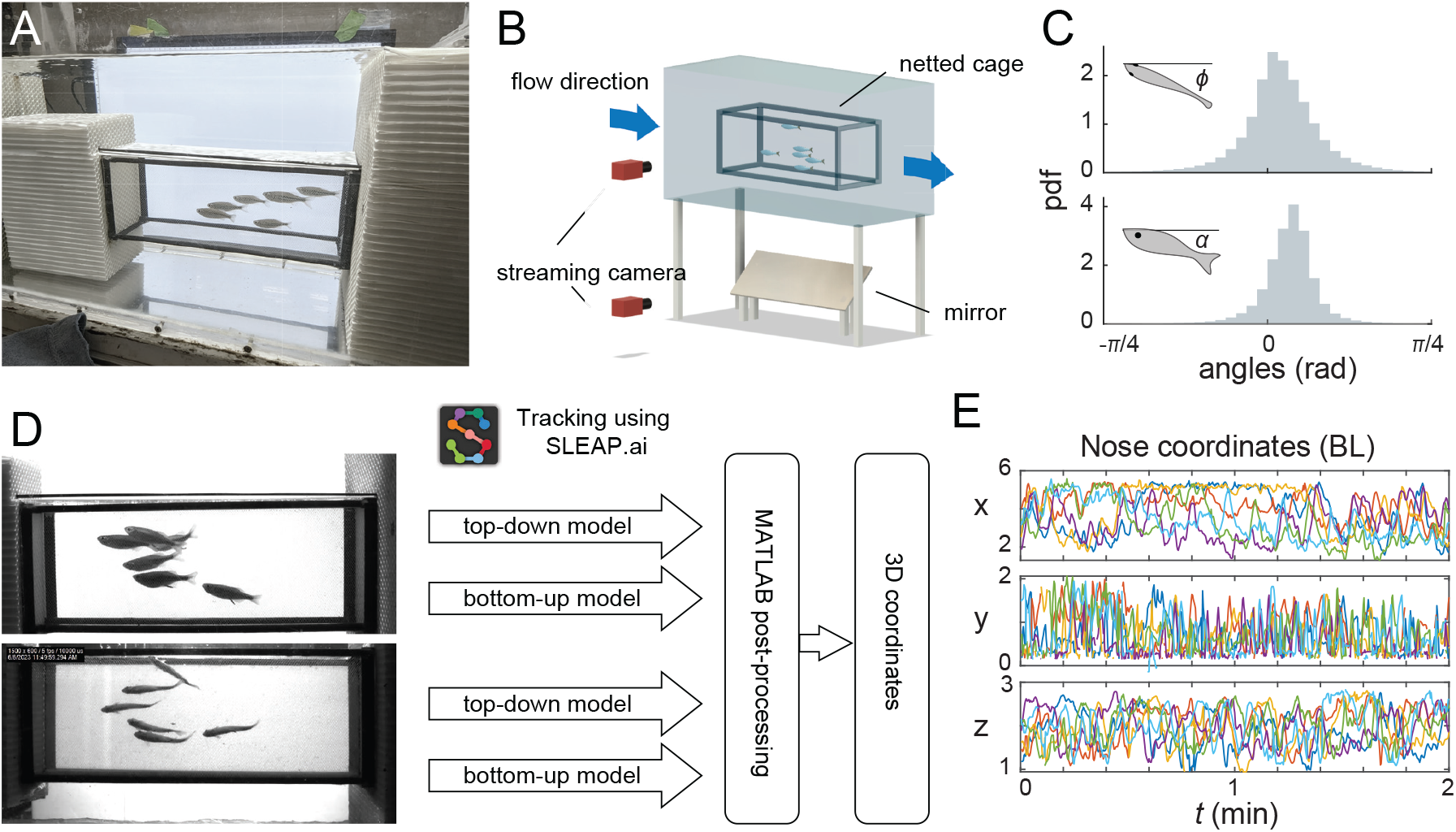
A machine vision pipeline for tracking fish schools in 3D. (A) A photo and (B) a schematic of the experimental setup. Note that the danio schools swam within a mesh chamber suspended within a recirculating flow tank that greatly reduced boundary layer effects (see Methods and Supplemental Material). (C) Histograms of both yaw *ϕ* and pitch *α*, centering around zero, i.e. aligning with the flow. (D) A diagram of the video-processing pipeline. (E) a two-minute sequence of the coordinates for the nose of all six fish within a school. Different colors represent different individuals.

### 2.1 Giant danio schools are transient and never planar

We begin by demonstrating the behavior of the entire school. Fig. 3A shows a time series of fish speed. Calculated as the mean speed of all fish relative to the chamber (in the lab frame), this metric characterized the school’s activity in the chamber. In the first hour, fish moved relatively quickly at approximately 1/4 BL/s. Then, their speed gradually decreased until it reached a baseline of around 0.15 BL/s. This suggests that it takes giant danio schools an hour to adapt their behavior to the flow. Thus, we limit our following analysis to the data after the first hour. The school’s vertical structure also exhibited an initial transient period (Fig. 3B). We measured school height as the vertical separation between the noses of the highest and lowest fish. Initially, the school maintained a relatively flat formation about one body-length tall, approximately 4.8 times the average height of an individual fish. After the first hour, school height fluctuated around a mean of 1.2 body-lengths, or 5.5 times fish body height (N=324,000). Notably, the fish school was never planar: less than 0.5% of all frames showed a school flatter than 0.25 body-length.

**Figure 3.**
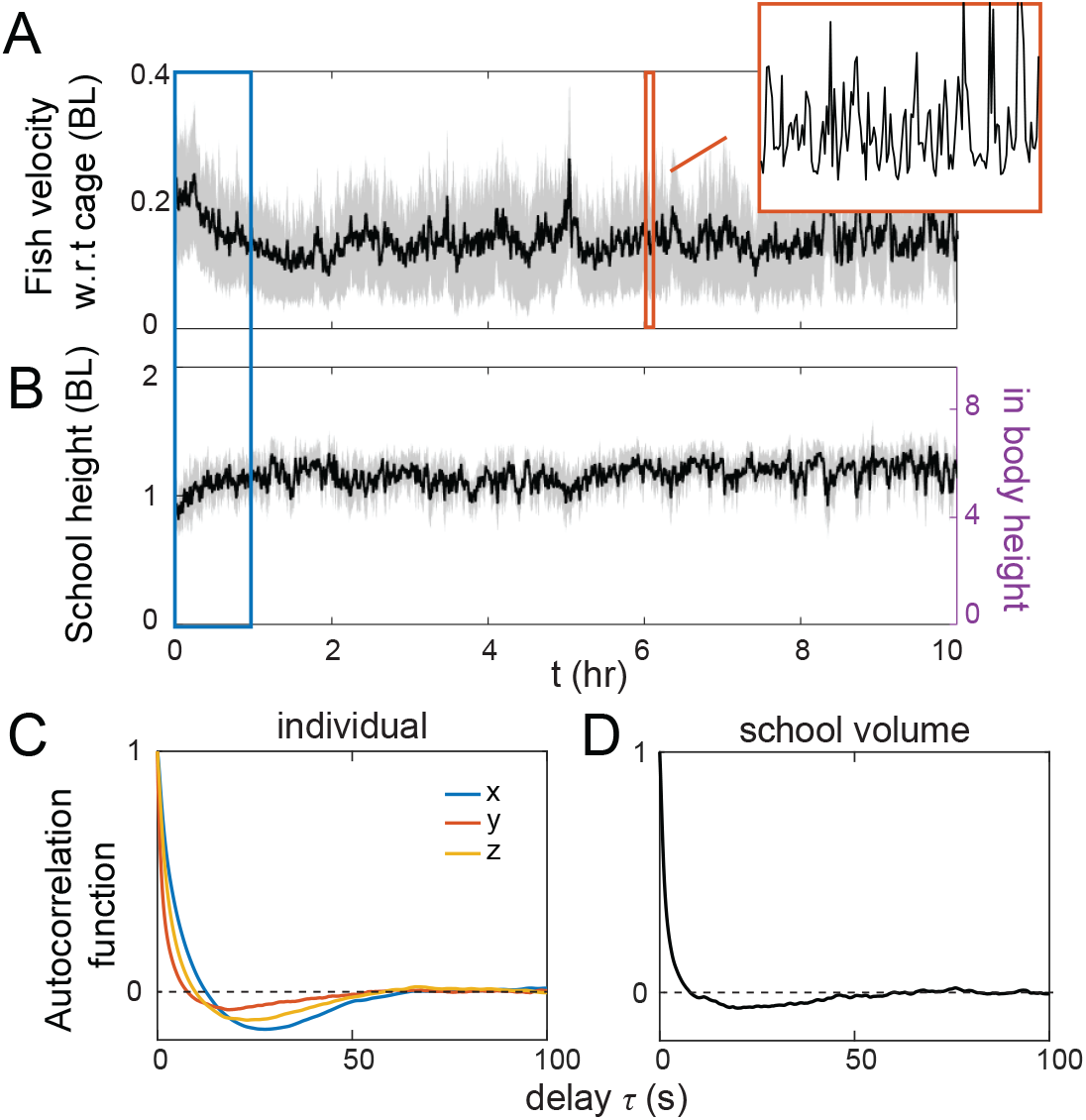
Fish schools are three-dimensional and transient. Time series of the (A) average fish speed with respect to the chamber, and (B) height of the school. Black lines represent moving averages and the gray regions represent standard deviations across 1-min-long moving windows. The transient period is marked in blue. (C-D) Autocorrelation functions for (C) individual fish coordinates and (D) school volume.

Both time series (Fig. 3A-B) also demonstrate that giant danio schools are in constant motion, actively rearranging their relative positions. To characterize this transient behavior, we calculated the autocorrelation function ACF(*τ*), which captures how a variable measured at two time instants *τ* apart correlates with each other (see Methods for details). This function typically decays from 1 at *τ* = 0 (perfect correlation with itself) to 0 when *τ* is so large that the two values are no longer correlated. The ACF for individual fish’s coordinates and the school’s volume indeed decayed with *τ* (Figure 1D-E). The de-correlation timescale was around 50 seconds for individual fish and around 10 seconds for the whole school. The school’s timescale was much shorter than the individual’s because the movement of any of the six fish in the school changes the school’s volume.

### 2.2 Fish schools rarely form a diamond

We used our data to directly test the conjecture posed by Weihs and Lighthill [3, 4]. This required a mathematical definition of the formation. We considered a fish quadruple to be in a diamond formation if they satisfied the following conditions (Fig. 4 A):

**Figure 4.**
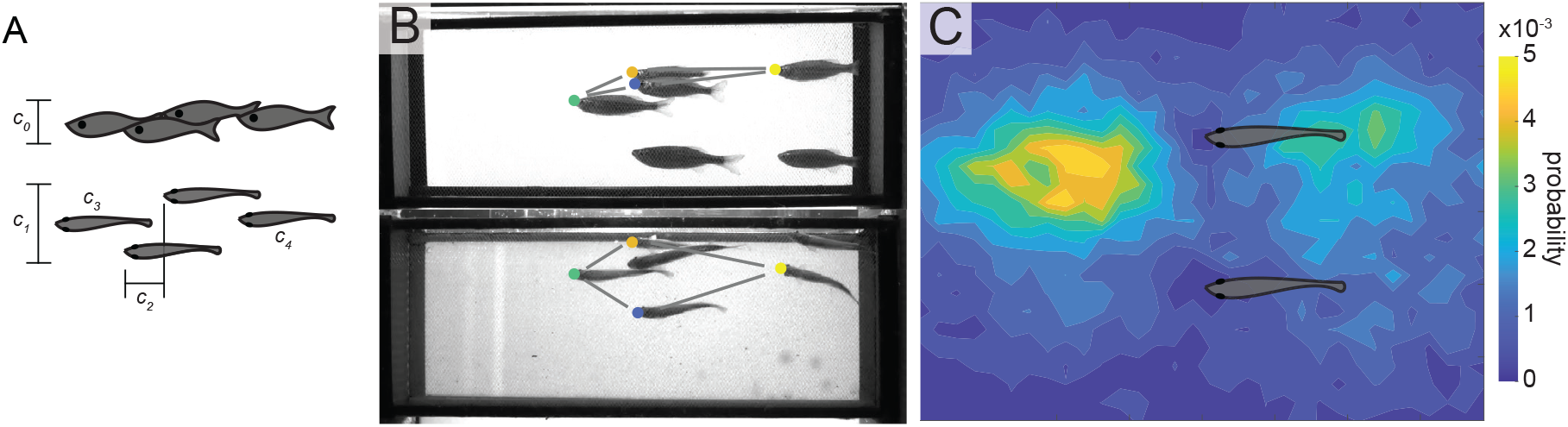
Fish rarely adopt diamond formations. (A) Diamond formations require a quadruple of fish satisfying conditions relating to school height (*c*_0_), school width (*c*_1_), streamwise separation between the fish pair on the outside (*c*_2_), and the presence of a leader and a follower (*c*_4_). (B) an example of the diamond formation identified by the automated algorithm. Such frames are rare in our recordings.

- The quadruple’s height *c*_0_ < 0.25 BL;
- Its width *c*_1_ < 1 BL;
- There exists a pair of fish side by side with streamwise separation *c*_2_ < 0.25 BL;
- There exists a leader *c*_3_ and a follower *c*_4_ in front of and behind the pair.

We designed an algorithm to automatically detect frames with diamond formations in our videos. Fig. 4B shows a representative frame identified by the algorithm. A quadruple of fish in this frame closely resembles the diamond formation proposed by Weihs and Lighthill [3, 4]. However, such frames were extremely rare in our experiments. Our video data provided 264,173 frames out of 324,000 total frames with at least four fish in the same plane. Of these, less than 0.1 % (262) were diamond formations. Fig. 4C shows the probability distribution of all fish around a side-by-side pair, i.e. satisfying conditions *c*_0_ − *c*_2_. The distribution of fish is almost uniform except for the region in front of the pair. Indeed, in our search, 1.5 % of frames with four fish in a plane had a side-by-side pair and at least a leader in front. This probability is much higher than having a fish trailing the pair, at 0.8 %. The hydrodynamic benefit proposed by the diamond model requires a minimal school with at least a pair of side-by-side fish and a trailing fish. We discovered that such a scenario is extremely rare for schools of giant danios under our experimental test conditions.

### 2.3 Fish most frequently adopt ladder formations in 3D

Although we never observed fish in schools maintaining any one formation for a prolonged period, a preferred unique ladder-like 3D arrangement was identified. Fig. 5AB show the probability maps of finding a neighbor around a fish located in the middle of the plot. Bright spots indicate the most likely location of a neighbor (N=2,482,308). Fig. 2A shows that a fish is likely to find a neighbor half a body-length either in front of it or behind it. The side view of this 3D heatmap (Fig. 5B) further revealed that the neighbor is likely below if in front, and above if behind. This unique ladder-like 3D arrangement can also be observed in Fig. 1C and 2A. To the knowledge of the authors, this ladder formation has never been reported.

**Figure 5.**
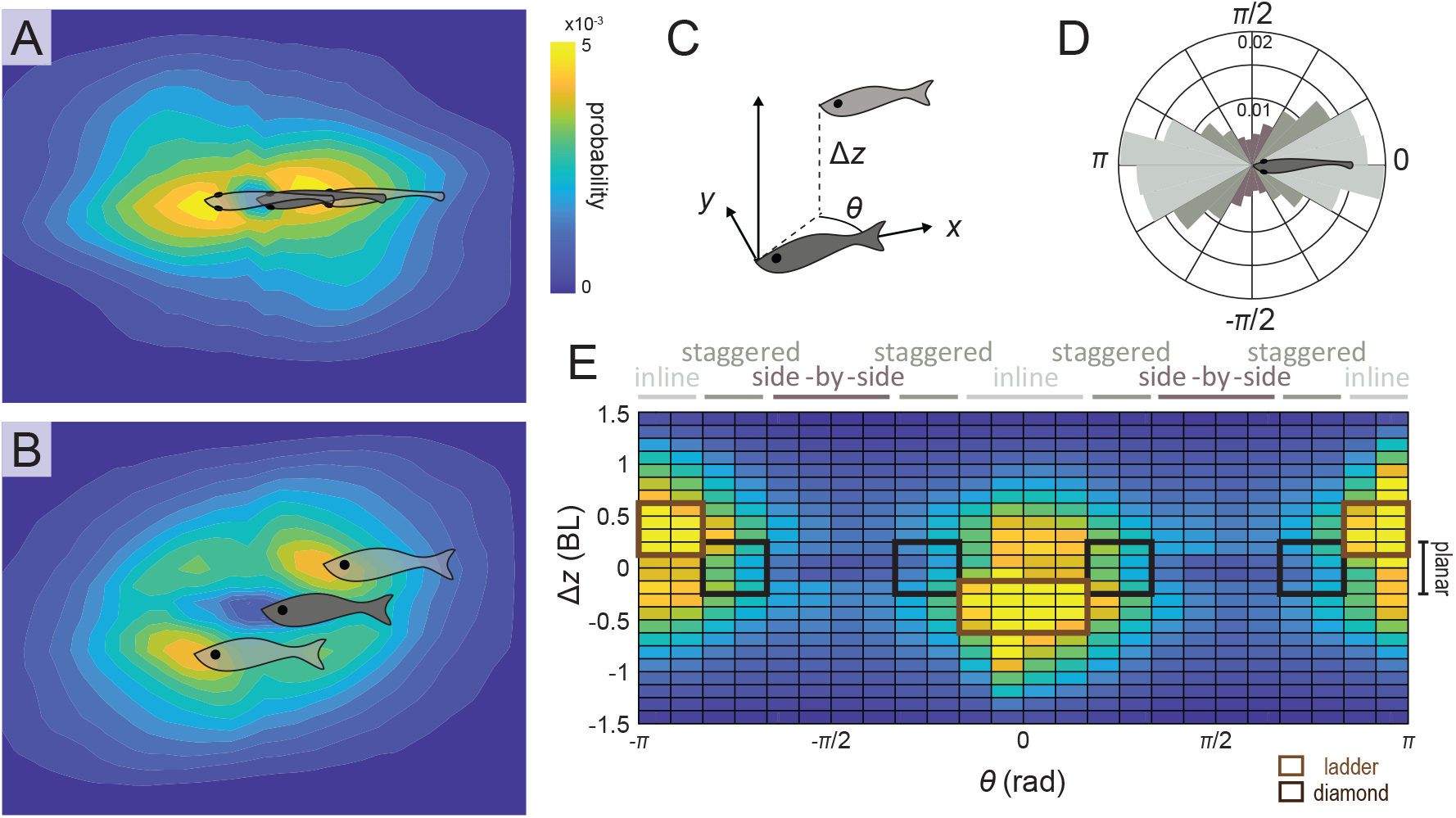
Fish pairs prefer vertically separated ladder formations. (A) A bottom view and (B) a side view of the probability distributions of finding a neighbor next to a central fish fixed at the center of the heat map. (C) A cylindrical coordinate representation of a fish pair’s formation. (D) A probability distribution of a neighbor’s bearing angle *θ* for all planar pairs. (E) A full probability distribution of a neighbor’s *θ* and *z*. Brown boxes denote ladder formations while black boxes denote planar staggered formations that would be prevalent in diamond formations.

The probability of finding a neighbor within a cylindrical zone surrounding an individual fish was calculated (Fig. 5C). A neighbor’s relative location can be characterized by the bearing angle *θ* and (C) The probability distribution of other fish next to a side-by-side fish pair. vertical separation *z*. The relative bearing angle *θ* between a fish pair defines whether their formation is inline, staggered, or side-by-side while their vertical separation *z* determines whether they are in a plane (see methods for more detail). Fig. 5D shows the distribution of *θ* for all planar pairs (N=62,212). 54.1% of these fish pairs were in inline formation, with a *θ* close to 0 or *pi*. Staggered formation was the second most likely at 39.5% and side-by-side formation was the least likely at 15.5%. Overall, planar formations (|*z*| < 0.25 BL) only consisted 25.1% of all fish pairs. This result suggests that the probability for a whole school of six giant danios to be in a plane is extremely low, around (25.1%)^5^ < 0.01%. This explains why the school height remained significant in Fig. 3B.

The full heat map (Fig. 5E) reveals how most pairwise formations are offset in the vertical direction *z*. The most probable locations of a neighbor occur when *θ* is around *π* or −*π* and Δ*z* around 0.5 BL, and when *θ* is around 0 and Δ*z* around -0.5 BL. These locations correspond to the ladder formation shown in Fig. 5AB. Summing all the probabilities within the black box, ladder formations accounted for 79.0% of all formations. In contrast, only 7.6 % of all fish pairs were in planar staggered formations, a configuration where fish in diamond formations would occur. Thus, Fig. 5E again demonstrates that diamond formation rarely occurs in giant danio schools.

### 2.4 Fish schools polarize more and elongate in a faster flow

We characterized how fish within schools adjust their formation as average water velocity increased from 1.6 to 5.6 BL/s. We limited the duration at each flow speed to five minutes and excluded the first minute in our analysis. While the school as a collective was still actively adjusting to flow in this period (Fig. 3AB), it only took several seconds for individual fish to adapt their swimming speeds to the increased incoming flow.

Fish pairs at all speeds adopted vertically separated formations similar to those in our 10-hour experiments at 2.4 BL/s (Fig. 4). Fig. 6A shows a heatmap from the side view for schools at three representative speeds, 1.6, 3.2, and 4.8 BL/s. At the lowest speed, a majority of the fish pairs were stacked vertically. As flow speed increased, the fish on top retreated and the formation became sheared. As a result, the school elongated as the flow rate increased. This elongation is statistically significant (p<0.001) (Fig. 6). This adaptive response to flow is similar to that observed for other collective systems such as the rafts of fire ants *Solenopsis invicta* [17].

**Figure 6.**
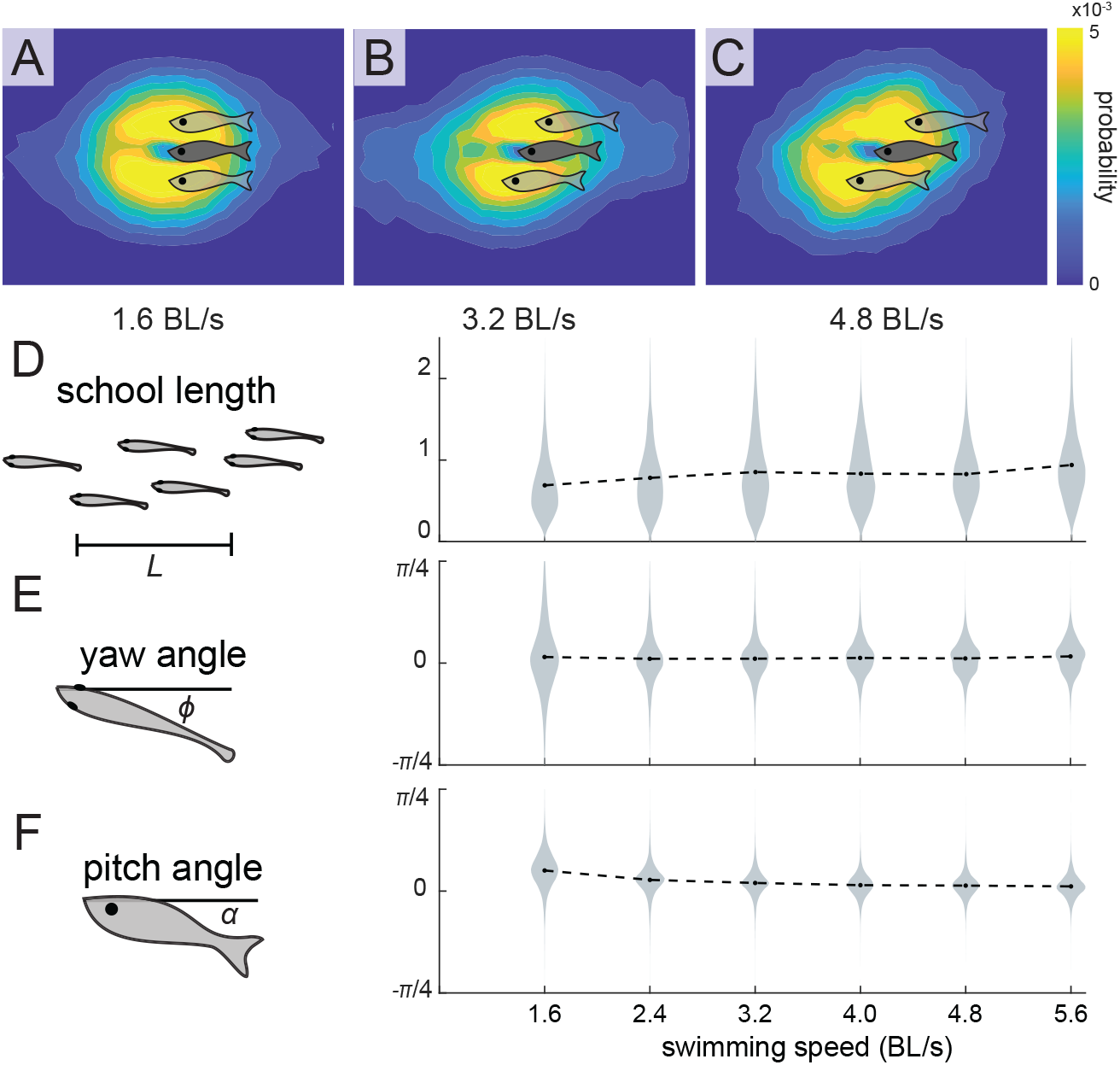
Fish schools become more polarized and more elongated in faster flow. (A-C) Heatmaps from the side view when the school swims at (A) 1.6 BL/s, (B) 3.2 BL/s, and (C) 4.8 BL/s. (D) school length *L*, (E) yaw *ϕ*, and (F) pitch *α* as a function of swimming speeds.

Fig. 6C-D show that while schools are polarized across all speeds, the polarization is stronger at faster speeds. The variances in yaw *ϕ* and pitch angles *α* with respect to the flow decrease with the flow speed. Note that the mean of the pitch angle *α* is slightly positive at low speed. This pitching behavior at low swimming speed has been previously described in leopard sharks (*Triakis Semifasciata*)[18], clearnose skate (*Raja eglanteria*)[19], rainbow trout (*Oncorhynchus mykiss*)[19], and giant danio [2], and been hypothesized to allow fish to generate lift and maintain vertical position. Our results indicate that the same mechanism may be used in schooling giant danio.

## 3 Discussion

Here we present a 10-hour three-dimensional characterization of fish school formations. By tracking polarized schools of giant danios swimming against flow in an experimental laboratory setting, we reveal that only 25.2 % of all fish pairs are in planar formations. This includes 13.6% in inline formations, 7.6 % in staggered formations, and 3.9 % in side-by-side formations (Fig. 7). The canonical diamond formation, widely cited as a preferred position for energy saving by fish within schools [3, 4, 20], was present less than 0.1 % of the time. A significant number of fish pairs are vertically separated by more than half a body-length apart. Notably, a “ladder formation” emerged as the most probable for schooling giant danios, appearing in 79.0 % of all fish pairs.

**Figure 7.**
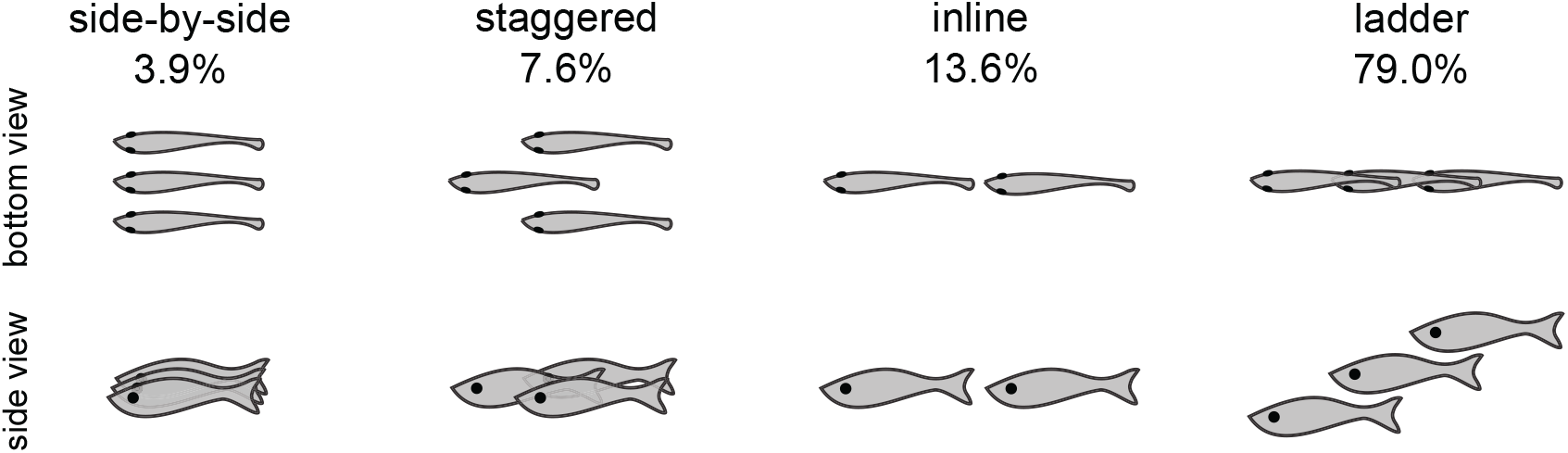
A diversity of pairwise formations and their probability identified in this study

To explore whether this ladder formation allows fish to gain hydrodynamic benefits, it is instructive to revisit the original hydrodynamic arguments for diamond formations [3, 4, 20]. Weihs argued that schooling fish would benefit from being in a diamond formation due to two hydrodynamic effects, one along the swimming direction (streamwise) and another perpendicular to the swimming direction (spanwise) [3]. The streamwise effect considers the fact that on average, a free-swimming fish leaves a jet behind going backwards. Thus, it is energetically unfavorable for a follower to be directly behind another fish. While fish confined in shallow water have to stagger horizontally to obtain this hydrodynamic benefit, when allowed to swim freely in all three-dimensions, they can also stagger in the vertical direction, resulting in the ladder formation.

The second proposed hydrodynamic benefit that fish in diamond formations could enjoy requires significant coordination among individuals. Weihs argued that if fish pairs that are side-by-side with each other beat their tails *anti-phase*, by symmetry, it would be as if they were beating their tails against a wall nearby, and therefore boosting their thrust [3]. In reality, this requirement may be difficult to satisfy. Coordinating tail-beats among individuals could be challenging, especially when fish are within large schools with hundreds or thousands of individuals. Furthermore, the observation that goldfish match each other’s vortex phase predicts that a side-by-side fish pair would in fact beat their tails *in phase* [21]. This suggests that the second benefit for diamond formations is perhaps seldom achieved and that the ladder formation may be just as efficient as the diamond formation.

A key result of this study is the finding that fish within a school are constantly rearranging their relative positions and do not maintain any one position for long periods of time (Fig. 7). This result, when combined with previous direct metabolic measurements of schooling energetics in this same danio species [2], raises the question of how fish moving around into different positions could reduce their cost of locomotion compared to an individual swimming alone. Under similar experimental conditions, giant danio in schools reduced their cost of transport and energy expenditure at higher speeds compared to solitary individuals [2]. Notably, the energy expenditure per tail beat was reduced by as much as 56% compared to solitary swimming [2]. Overall, fish schools act as a hydrodynamic shelter for the individuals within, as demonstrated by our previous metabolic study of danio schools swimming in turbulence [22]: the school sheltered individuals from the hydrodynamic perturbations in turbulence and significantly lowered the energetic cost of locomotion compared to solitary swimming in turbulent fluid environment.

The direct measurement of energetic saving by collective motion in this same species [2, 22] raises the question: how could fish reduce their locomotive energy expenditure in a school when they are frequently changing their relative positions? The resolution of this question lies in the diverse array of formations that offer energy-saving benefits. Numerical simulations and experimental work—involving live fish, flapping hydrofoils, and robotic models—have revealed that energy savings occur across a wide spectrum of relative positions, and the classical diamond formation may not be optimal [23, 24, 25]. Fish are predicted to reduce their cost of transport compared to solitary swimming when swimming in inline formations [26, 27, 28, 29, 30], side-by-side formations [31, 32], staggered formations [33, 34, 35], rectangular formations [20, 36], and vertically aligned formations [37]. This diversity of energy-saving positions suggests that fish schooling behavior is highly permissive in allowing hydrodynamic energetic reduction in swimming cost even though fish in a school are in constant motion relative to each other, as demonstrated here. The previously measured energy reduction could result from the accumulation of all savings retained under each of the formations occupied.

The hydrodynamic interactions between fish in three-dimensional formations remain an open question currently under investigation. Previous research has primarily focused on two-dimensional formations, leaving several critical questions unanswered. How does the fluid field develop and propagate around fish in ladder formations? Is energy saving enhanced when tail beats are synchronized across different formations? Future studies employing numerical simulations, hydrofoil experiments, and swarm robotics promise to provide insights into new classes of three-dimensional fish formations [5]. These approaches will significantly improved our understanding of the dynamics of the canonical collective behavior represented by fish schooling.

Three-dimensional tracking of animal collectives continues to yield exciting and fruitful discoveries, offering unprecedented insights into complex biological systems. Our computer vision pipeline contributes to this rapidly evolving landscape, complementing recent advancements such as the tracking of fireflies in their native habitat, which has unveiled crucial self-organizing mechanisms [38, 39], and the 3D tracking of brine shrimp movements, which has illuminated the hydrodynamic consequences of collective migration [40]. Future applications may extend to tracking fish schools in the wild and swarms of other aquatic animals, advancing our understanding of collective behavior across species and scales.

## 4 Materials and Methods

### 4.1 Fish care

In our experiments, we used giant danios (*Devario aequipinnatus*) that were 6.0 ± 0.6 cm long, 0.7 ± 0.1 cm wide, and 1.3 ± 0.1 cm tall. They were acquired from a local commercial supplier near Boston, Massachusetts, United States of America, and housed in a 37.9 L aquarium. The aquarium was equipped with a filtration system and was temperature and aeration controlled (at 28°C and >95% air saturation). Water changes (up to 50% exchange ratio) were carried out weekly. Fish were fed ad libitum daily (TetraMin, Germany). Animal holding and experimental procedures were approved by the Harvard Animal Care IACUC Committee (protocol number 20-03-3).

### 4.2 Experiments

Experiments were conducted in June 2023. A water channel (overall test channel 50 cm long, 30 cm wide, and 30 cm tall) was used to provide constant flow to the fish schools to encourage active directional schooling behavior and polarized formations. Schools were placed within a rectangular cage made with carbon fiber rods and nylon netting to form walls (30 cm × 11.5 × 11 cm) located within the test section of the water channel (Fig. 2). Without this cage, fish preferred staying in regions of slower flow next to the walls of the water channel, taking advantage of the boundary layer that develops at the walls. The cage greatly reduced the wall effect, and flow within the cage was much more uniform, as verified by PIV (Particle Image Velocimetry) calibration (Fig. S1). Flow velocity at the net walls of the chamber was only less than 28% slower than the free stream velocity at the center (Fig. S1). In contrast, without the cage, flow velocity would have decreased to zero at the boundaries, i.e. a 100% reduction in speed. This substantial improvement in flow uniformity mitigated the boundary layer effects on fish school behavior and reduced the frequency with which fish were located near the boundaries.

The experiments were recorded using a pair of streaming cameras AOS PROMON U1000 fitted with aspherical lenses (4-12 mm). Videos were taken from the side and the bottom at 5 fps. Both views were back-lit. We placed an LED panel (HUION A3) at the back of the flow channel for the side view, and two diffused LED spotlights (Nila Zaila) above the water channel for the bottom view. A transparent acrylic plate was placed on the water surface to avoid shadows caused by surface distortion. Videos were streamed to a hard drive (500G) using AOS Imaging Studio. All frames were cropped from 1920×1080 to 1500×600 while being saved. Each 10-hour video took up 2.5 GB in raw format and 32 MB after being converted to mp4 format.

The night before each experiment, we transferred six giant danios into the net cage to allow for at least 12 hours of acclimation. At the beginning of each experiment, we applied gentle flow to clear away debris in the water channel. Two experimental protocols were used in this study. In the steady flow experiments, we provided a constant flow at approximately 14 cm/s or 2.4 BL/s. Throughout the 10 hour experiment, all six fish swam continuously and the entire experiment was shrouded to avoid external stimuli affecting fish schooling behavior. Recording started 15 minutes after the flow started. In separate ramping flow experiments, we increased flow velocity by 0.8 BL/s every ten minutes, varying from BL/s to BL/s. We performed six such trials, separated at least an hour apart for fish to rest in static water in between replicate trials.

### 4.3 Fish tracking

We used SLEAP.ai, a deep-learning-based multi-animal markerless tracking algorithm [16]. We manually labeled the nose and peduncle (the narrowed region before the caudal fin) in approximately 150 frames in both views. Two tracking methods were used: a top-down method used a network to find the location of each fish first and then used a separate network to locate its nose and peduncle (the area near the base of the tail). A bottom-up method used a network to find all the noses and peduncles in a frame before assigning them to each fish. Neither method provided perfect tracking for our videos but one method often succeeded where the other one failed. Therefore, we used both methods and designed a post-processing script in MATLAB to improve tracking outcome. See supplemental materials for a pseudo code. In this automated script, we kept the coordinates if they were predicted by both methods. If a coordinate was only identified in one method but not the other, we decided whether to remove it based on a cost function that considered both the fish’s location in the previous frame, and the fish’s streamwise locations (x-coordinate) in the other view. The script then combined the processed information to calculate the 3D trajectories of each fish. Note that as fish cross paths with each other, their identity may be swapped. Therefore, our method cannot yet answer questions such as how often fish switch leadership. The training of all neural nets was performed on Google Colab, and the inference was performed on the Tiger Cluster of Princeton University. On a node with one GPU, four CPUs, and 120 GB memory, tracking all 180,000 frames in a 10-hour video using each method and from each view took around 2.5 to 3 hours. Post-processing took another 5 hours on a node with 1 GPU, 4 CPUs, and 64 GB memory.

### 4.4 Pairwise formations

In this manuscript, fish pairwise formations were defined based on the relative positions between each fish pair within the school. A fish pair is considered to be planar if their relative displacement in the vertical direction is within 0.25 body-length, approximately 1.25 times the height of a fish. Following common practice, we define all staggered formations, inline formations, and side-by-side formations as planar formations. A planar (or at least near-planar) orientation is necessary for fish to interact hydrodynamically based on current hypotheses and models considered further in the Discussion. These formations are distinguished based on the bearing angle, *θ* (Fig. 5C). 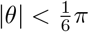 (neighbor in the front) and 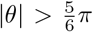 (neighbor in the back) were defined as inline formations. 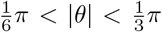 (neighbor at an angle to the front) and 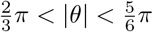 (neighbor at an angle to the back) were defined as diamond formations. 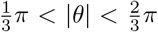 were defined as side-by-side formations. The emergent ladder formations were defined based on the brown box in Fig. 5E: for −0.625 BL < Δ*z* < −0.125 BL when 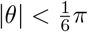, and for 0.125 BL < Δ*z* < 0.625 BL when 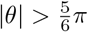. Based on this definition, if all fish pairs were randomly distributed, staggered, inline, and side-by-side formations would take up 1/3 of planar formations, and all formations including ladder formations would be equally probable at 1/18 or 5.6%.

For the calculation of the probabilities of pairwise formations, we removed the initial transient periods (1 hr) as shown in Fig 3AB. In addition, we excluded individuals immediately next to our netted cage to avoid the bias introduced by the boundaries. Only fish pairs that are within 4.0 bodylengths apart radially, and 1.5 body-lengths apart vertically were included. Fish pairs outside of this range likely have negligible hydrodynamic and social interactions with each other. The total number of fish pairs that satisfied these criteria was used as the denominator for all probabilities reported in this manuscript.

### 4.5 Fish school metrics

In this project, we calculated fish speed, school height and school length. Fish speed (Fig. 3A) was calculated as the average speed of all six fish relative to their arena, i,e, the netted cage. School height (Fig. 3B) was calculated as the vertical gap between the highest and the lowest fish detected in the frame. Since the algorithm failed to identify all six fish in some frames, the school height calculated was a lower bound. Any undetected fish would only increase the value, further showcasing that danio schools seldom from planar formations. Lastly, the school length reported in Fig. 6D was calculated as the streamwise distance between the second fish and the second last fish. This was to avoid situations where there was a solitary fish far away, either in front of, or behind the school.

To characterize the persistence timescale for fish coordinates *x*(*t*), *y*(*t*), *z*(*t*), and the school volume *V* (*t*), we calculated autocorrelation functions (ACF) of these time series (Fig. 3). An autocorrelation function calculates how a variable de-correlates with itself given the delay *τ* between the two measurements. It was calculated using the formula 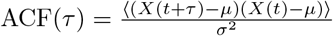. Here, *X*(*t*) is a shorthand for any of the four time series to be calculated, and *μ* and *σ* are the time series’ mean and standard deviation across a 2-min-wide moving windows. ⟨…⟩ represents an average across all individuals and all frames. By definition, ACF=0 at *τ* = 0. As the delay between two measurements of the same variable increases, *τ* → ∞, *X*(*t* + *τ*) and *X*(*t*) become un-correlated, and ACF approaches zero.

Finally, we examined whether the school length, fish’s yaw, and fish pitch correlates with the swimming speed using a general linear model (GLM, Fixed factors: water velocity) followed by Tukey HSD posthoc tests. The test was implemented in SPSS (v29.0.0, SPSS Inc, Chicago, IL, USA).

## Acknowledgement

We would like to acknowledge the support from ONR MURI grant N00014-22-1-2616 (co-PIs R. Nagpal and G.V. Lauder) and the The Company of Biologists Travelling Fellowship (H.Ko). In addition, H.Ko is supported by the James S. McDonnell Foundation’s Postdoctoral Fellowship for Understanding Dynamic & Multi-scale Systems. Y. Zhang is supported by a Postdoctoral Fellowship of the Natural Sciences and Engineering Research Council of Canada (NSERC PDF - 557785 – 2021) and a Banting Postdoctoral Fellowship (202309BPF-510048-BNE-295921) of NSERC & CIHR (Canadian Institutes of Health Research). The computation presented in this article were performed on computational resources managed and supported by Princeton Research Computing.

